# Chromosomal fusion drives sex chromosome evolution in treehoppers despite long-term X chromosome conservation

**DOI:** 10.1101/2024.07.12.603334

**Authors:** Daniela H Palmer Droguett, Micah Fletcher, Sarah Kocher, Diogo C Cabral-de-Mello, Alison E Wright

**Affiliations:** Ecology and Evolutionary Biology, School of Biosciences, The University of Sheffield, Sheffield, UK; Ecology, Evolution, and Behavior Program, Michigan State University, East Lansing, MI, USA; Department of Biology, The University of Texas at Arlington, Arlington, TX, USA; Department of Ecology & Evolutionary Biology and the Lewis-Sigler Institute for Integrative Genomics, Princeton University, Princeton, NJ, USA; Howard Hughes Medical Institute, Princeton University, Princeton, NJ, USA; Department of General and Biology, São Paulo State University (UNESP), Institute of Biosciences, Rio Claro

**Keywords:** sex chromosomes, neo-sex chromosomes, chromosomal fusion, chromosomal rearrangements, karyotype evolution

## Abstract

Sex chromosomes follow distinct evolutionary trajectories compared to the rest of the genome. In many cases, sex chromosomes (X and Y, or Z and W) significantly differentiate from one another resulting in heteromorphic sex chromosome systems. Such heteromorphic systems are thought to act as an evolutionary trap that prevents subsequent turnover of the sex chromosome system. For old, degenerated sex chromosome systems in which turnover is unlikely, chromosomal fusion with an autosome may be one way that sex chromosomes can ‘refresh’ their sequence content. We investigated these dynamics using treehoppers (hemipteran insects of the family Membracidae), which ancestrally have XX/X0 sex chromosomes. We assembled the first chromosome-level treehopper genome from *Umbonia crassicornis* and employed comparative genomic analyses of 12 additional treehopper species to analyze X chromosome variation across different evolutionary timescales. We find that the X chromosome is largely conserved, with one exception being an X-autosome fusion in *Calloconophora caliginosa*. We also compare the ancestral treehopper X with other X chromosomes in Auchenorrhyncha (the clade containing treehoppers, leafhoppers, spittlebugs, cicadas, and planthoppers), revealing X conservation across more than 300 million years. These findings shed light on chromosomal evolution dynamics in treehoppers and the role of chromosomal rearrangements in sex chromosome evolution.

**Significance:** The evolutionary forces underlying sex chromosome stability versus turnover have been challenging to disentangle. We present the first chromosome-level treehopper genome and find evidence of long-term X chromosome conservation within treehoppers – and among treehoppers and other hemipteran insects. A key exception is the evolution of neo-XX/XY sex chromosomes via an X-autosome fusion. Sex chromosome-autosome fusions may play an important role in the evolution of otherwise ‘trapped’ (i.e., old and degenerated) sex chromosome systems.

## Introduction

In many clades, pairs of sex chromosomes are often observed to follow distinct evolutionary trajectories in which the X (or Z) chromosome remains functional and gene-rich while the Y (or W) undergoes functional degeneration and loss (Charlesworth, et al. 2005; Bachtrog, et al. 2011). The emergence of such heteromorphism is predicted to act as an evolutionary trap that impedes sex chromosomes from undergoing turnover to new genomic regions (Bull 1983; Bachtrog, et al. 2011), For example, evolutionary turnover from an XY to a ZW system can generate YY individuals, making such a transition unlikely if the Y is degenerated and YY individuals have low fitness. This idea been supported by several comparative analyses (Pokorná and Kratochvíl 2009, Gamble, et al. 2015, Nielsen et al. 2019) and anecdotally many clades with old sex chromosomes exhibit long-term sex chromosome conservation, such as birds, insects, mammals, and some groups of reptiles (Pokorná and Kratochvíl 2009; Nielsen, et al. 2019; Chauhan, et al. 2021; Li, et al. 2022; Toups and Vicoso 2023), however, there are notable exceptions (Vicoso and Bachtrog 2013; Pinto, et al. 2023). On the other hand, some clades maintain sex chromosome homomorphism which is thought to permit rapid and ongoing turnover in both sex chromosome identity and system (Gamble, et al. 2015; Jeffries, et al. 2018; Tennessen, et al. 2018; Balounova, et al. 2019; Darolti, et al. 2019). What degree of sex chromosome divergence is necessary to create a trap and whether this affects all chromosomes equally remains unclear.

We do know that key exceptions to the evolutionary trap model of sex chromosome evolution are driven by chromosomal fusions between sex chromosomes and autosomes (Maddison and Leduc-Robert 2013; Pennell, et al. 2015; Sigeman, et al. 2022; Castillo, et al. 2023). These fusions can generate new sex chromosome systems and ‘refresh’ the sequence content of the sex-linked genome. Chromosomal rearrangements can have important functional genomic impacts, either directly by changing the sequences of regulatory and protein coding regions (Stewart and Rogers 2019) or more indirectly by altering linkage and recombination among selected loci (Cicconardi, et al. 2021; Näsvall, et al. 2023). Fusions between an autosome and a sex chromosome present a special case with potentially more extensive impacts because of the resulting shift toward sex-linked inheritance of previously autosomal regions. These newly sex-linked regions will thus be subject to sex differences in selection, demography, and life history that can drastically alter their evolutionary trajectories (Rice 1984). Different evolutionary forces are predicted to drive fusions between sex chromosomes and autosomes, including sexually antagonistic selection and meiotic drive, however, the relative importance of each is unclear and appears to differ across taxonomic groups (Charlesworth and Charlesworth 1980; Pennell, et al. 2015; Anderson, et al. 2020). Identifying the structural genomic changes involved in such events and their evolutionary drivers are thus consequential for understanding how and why sex chromosomes evolve and persist.

Insects are a promising system for investigating the relative role of fusions in sex chromosome evolution. Karyotypic data indicates that many species exhibit frequent genome rearrangements, with significant changes in chromosome number (The Tree of Sex 2014; Blackmon, et al. 2017), and so we might expect autosome-sex chromosome fusions to be common if there was a selective advantage (Pennell, et al. 2015). While many insect species do exhibit neo-sex chromosomes (Yoshido, et al. 2011; Zhou and Bachtrog 2012; Zhou, et al. 2012; Maddison and Leduc-Robert 2013; Nguyen, et al. 2013; Vicoso and Bachtrog 2015; Blackmon, et al. 2017; Mongue, et al. 2017; Palacios-Gimenez, et al. 2018; Bracewell, et al. 2023; Decroly, et al. 2024), it remains unclear whether the frequency of fusions involving sex chromosomes mirrors genome wide patterns of rearrangement. There is growing evidence that the insect X chromosome has been conserved for long time periods (Chauhan, et al. 2021; Li, et al. 2022; Toups and Vicoso 2023), but the occurrence of neo-sex chromosomes appears to vary substantially across taxonomic groups (Mathers, et al. 2021; Bracewell, et al. 2023).

To investigate patterns of stability versus turnover of sex chromosomes, we employ whole-genome sequence data across 13 treehopper species spanning approximately 45 million years of evolution. Treehoppers (Membracidae) are a group of hemipteran insects best known for their morphologically diverse pronota or “helmets” (Prud’homme, et al. 2011; Fisher, et al. 2020). Knowledge of treehopper genome evolution has been relatively limited in comparison to other well-studied hemipteran groups such as aphids (Aphididae) (Jaquiéry, et al. 2012; Jaquiéry, et al. 2013; Li, et al. 2020; Mathers, et al. 2021) and planthoppers (Delphacidae) (Ma, et al. 2021; Ye, et al. 2021; Hu, et al. 2022), whose genomes show differing levels of synteny but X chromosome conservation within each clade. In contrast to aphids, treehoppers are obligately sexually reproducing and exhibit a variety of sex-related phenotypes like vibrational courtship signals and parental care. The most prevalent sex chromosome configuration in treehoppers based on karyotype information is XX/X0 (Halkka 1959, 1962, 1964; Halkka and Heinonen 1964), in which females carry two X chromosomes, and males carry a single X. However, multiple species carrying XX/XY systems have been identified by cytological studies (Kornhauser 1914; Tian and Yuan 1997; Anjos, et al. 2019), indicating the repeated emergence of new Y chromosomes. Cytological studies indicate that total chromosome number can range from 5 to 11 pairs of chromosomes, with the mode being 11 pairs. These data suggest that treehopper chromosomes undergo frequent chromosomal rearrangements.

Here, we build a new chromosome-level assembly for *Umbonia crassicornis*, generate genome assemblies using data from male and female individuals of 12 additional treehopper species spanning short, medium, and long-term evolutionary distances, and combine our data with published hemipteran genomes. We interrogate the relationship between chromosomal fusions and sex chromosome evolution and test the extent to which long-term conservation of the X chromosome previously observed across broad insect groups (Chauhan, et al. 2021; Li, et al. 2022; Toups and Vicoso 2023) is a feature of treehopper evolution.

## Results and Discussion

### Umbonia crassicornis genome assembly

We assembled the first chromosome-level treehopper genome from *Umbonia crassicornis* using 10x linked-reads and HiC data. The resulting genome assembly was 1.2 Gb. The average scaffold was 10,566kb and 70% of the genome was contained on the first 10 scaffolds of the assembly, whose sizes range from 54.3 Mb to 164Mb. These ten large scaffolds are coincident with the number of chromosomes observed by cytological analysis, as the species has 9 autosomal chromosomes plus an X element (Escudero and Virkki 1976). The ancestral diploid number for Membracidae is 10 autosomes + X0 (Emeljanov and Kirillova 1992; Kuznetsova and Aguin-Pombo 2015), implying that *Umbonia crassicornis* underwent a reduction in chromosome number caused by a fusion between two autosomes. Consistent with this cytological evidence, we observed that the largest scaffold (164Mb) of the assembly is roughly twice as large as the other nine major scaffolds (average 77.5Mb). We propose that the largest scaffold represents the product of an autosome-autosome fusion. The genome assembly is available on NCBI (PRJNA1122077).

### Identifying and comparing X-linked sequences across treehoppers

We next leveraged the *Umbonia crassicornis* genome assembly to identify the X chromosome in this species. We mapped paired-end reads from two male and two female samples to the genome assembly. We then calculated the log_2_ male to female coverage ratio for each of the ten major pseudo-chromosomes (hereafter referred to as chromosomes). For a species with an XX/X0 system, this approach is expected to yield a value of 0 (equal coverage among males and females) for autosomal sequences, and −1 (half the coverage in males compared to females) for X-linked sequences. Here we report results based on read mapping to the ten chromosomes. We repeated the analyses using the full genome assembly and results were qualitatively the same. Our results showed average log_2_ male to female coverage close to 0 for chromosomes 1 through 9, in contrast to chromosome 10 which showed an average log_2_ male to female coverage ratio of −0.978 (Figure 1). We further investigated sex differences in coverage in windows across each of the chromosomes to confirm the expected signatures for an XX/X0 system and observed a reduction of male coverage across the entire length of chromosome 10 (Figure S1). Based on this evidence we conclude that chromosome 10 is the X chromosome in *Umbonia crassicornis*.

**Figure 1.**
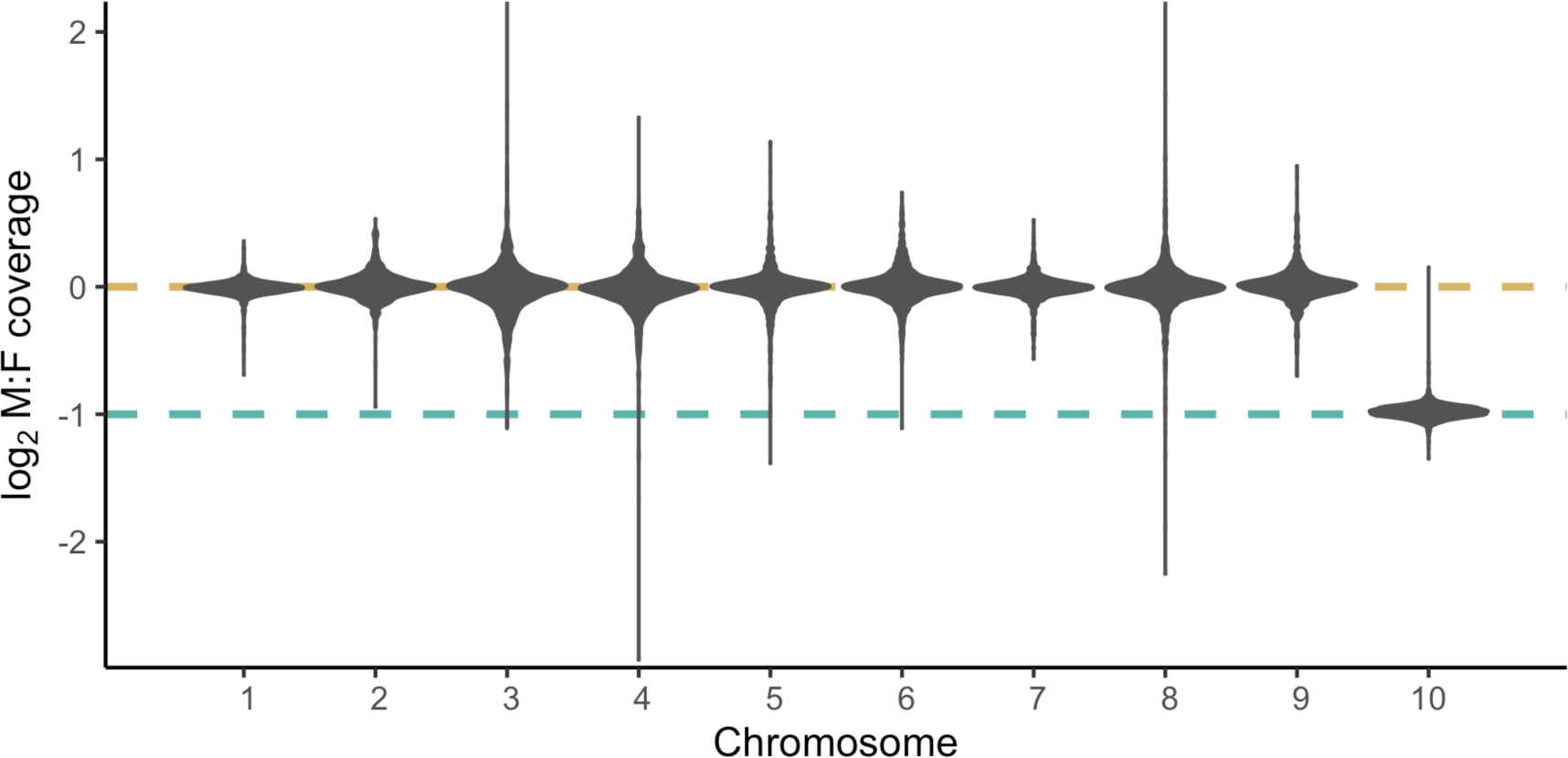
Sex differences in coverage across *Umbonia crassicornis* chromosomes show a reduction of male/female coverage for chromosome 10. Patterns of male/female sequencing coverage across the ten major scaffolds corresponding to the ten *U. crassicornis* chromosomes. Dashed lines in gold and teal show the expected coverage values for autosomal and X-linked regions, respectively. Chromosome 10 was classified as the X chromosome based on the halving of male/female coverage compared to chromosomes 1 through 9.

For each of the remaining 12 species (Figure 2a) we generated short-read sequence data for males and females (Table S1) and built de novo genome assemblies (Table S2). We then used the same coverage approach as above to identify autosomal and sex-linked sequences in each species. These species are known to vary in karyotype (Table S1), so comparing the repertoire of autosomal versus sex-linked sequences among species should reveal chromosomal rearrangements involving sex chromosomes (Lasne, et al. 2023). Since we did not have chromosome-level resolution for these other taxa, we anchored sequences back to the *Umbonia crassicornis* assembly to ask whether the identity of sex-linked sequences is shared. For all species but two, we found that most sequences with autosomal coverage patterns corresponded to autosomes in *Umbonia crassicornis* (Figure 2b). We also found that sequences with X-linked coverage patterns corresponded to the X chromosome in *Umbonia*, suggesting conservation of X chromosome identity across species, spanning 45 million years of treehopper evolution. From here, we refer to this shared X chromosome as the ancestral treehopper X chromosome.

**Figure 2.**
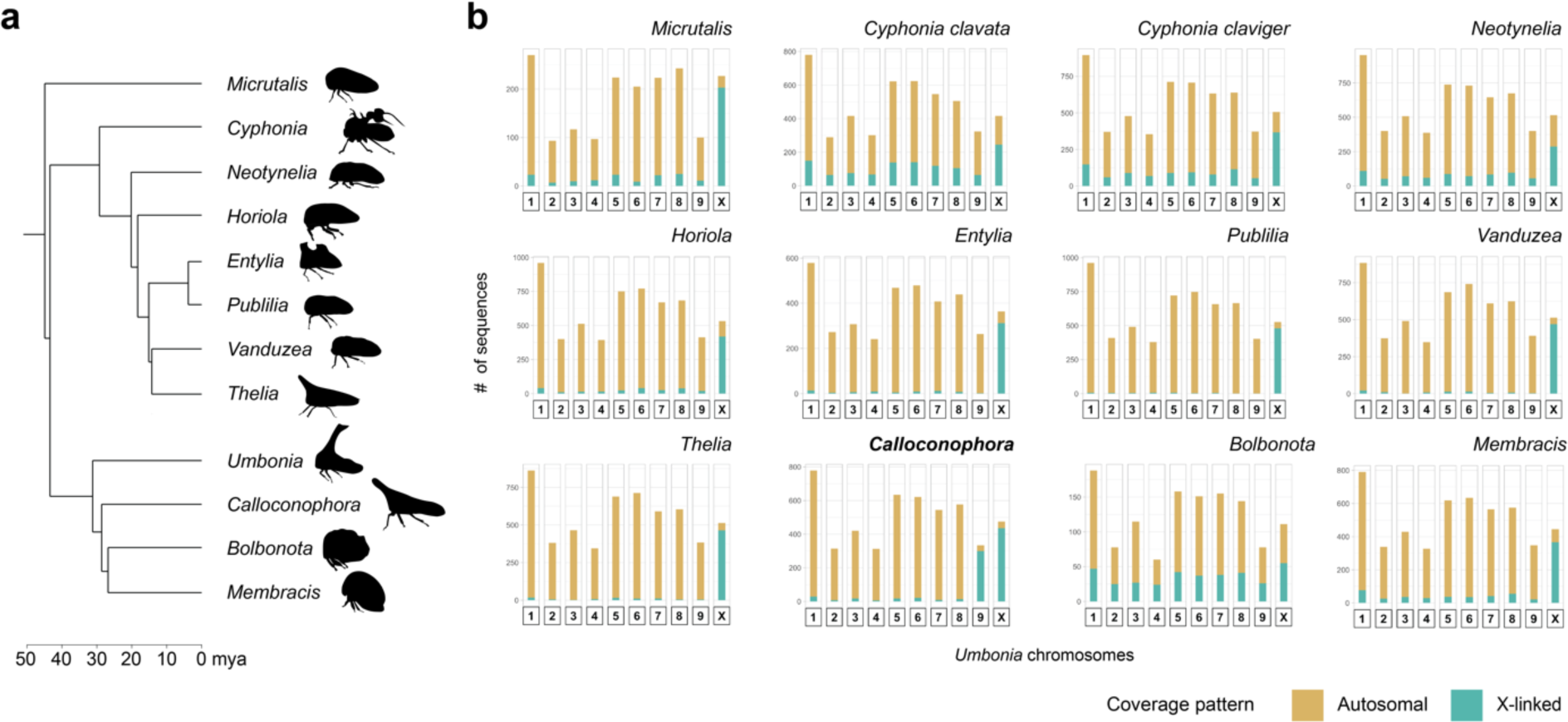
X chromosome identity is largely conserved across treehopper genera. (a) Phylogenetic tree of treehopper genera sampled for this study (adapted from (Fletcher 2023). (b) Pairwise comparisons of X- versus autosomal linkage between each focal taxon and *U. crassicornis*. Height of the bar indicates the number of focal taxon sequences that correspond to each *U. crassicornis* chromosome (along the x-axis), and color of the bar shows the number of sequences that are inferred to be autosomal (in gold) versus X-linked (in teal) in the focal taxon.

The only qualitative exceptions to this trend were *Calloconophora and Bolbonota*. Sequences with X-linked coverage patterns in *Bolbonota* were distributed evenly across *Umbonia* autosomes, although the greatest proportion were still found on the *Umbonia* X. This noise is likely a product of lower sequencing coverage for individuals of this species (Table S1). In contrast, *Calloconophora* showed clear X-linked coverage patterns for sequences corresponding to both the *Umbonia* X chromosome and chromosome 9 (which is autosomal in *Umbonia*) (Figure 2b). Given the frequent changes in chromosome number that are known to occur in treehoppers, we hypothesized that a chromosomal fusion occurred in *Calloconophora* between the ancestral sex chromosomes and the homolog(s) of *Umbonia* chromosome 9.

### Confirming a chromosomal fusion and neo-X in Calloconophora

To further investigate the presence of a sex chromosome-autosome fusion in *Calloconophora*, we used FISH to visualize the chromosomes and the location of telomeric sequences. This confirmed the presence of an X and a Y chromosome and revealed the X to be approximately 25% larger than the Y (Figure 3a). Notably, we observed signal corresponding to telomeric sequence toward the middle of the X chromosome, indicating the site of a chromosomal fusion between the ancestral X and an autosome (Figure 3a).

**Figure 3.**
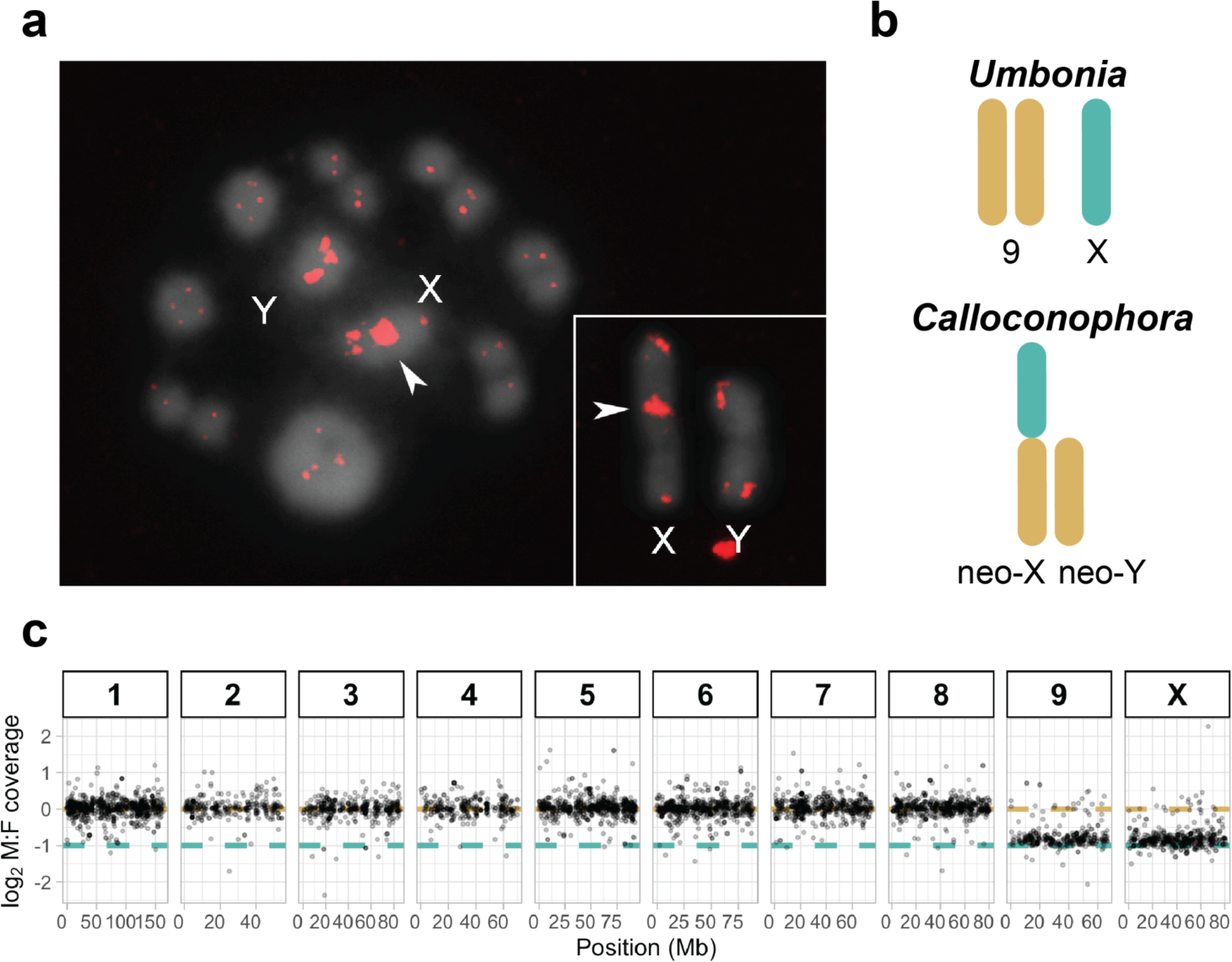
An X-autosome fusion underlies the formation of neo-sex chromosomes in *Calloconophora.* (a) FISH showing telomeric sequences in red at metaphase I (main panel) and mitotic metaphase (insert). Interstitial signal indicated by arrowheads shows the putative X-autosome fusion site. (b) Schematic of the inferred pre- (*Umbonia*) and post-fusion (*Calloconophora*) chromosomes involved in the transition from XX/X0 to XX/XY sex chromosomes. (c) Patterns of male/female sequencing coverage for *Calloconophora.* The numbered boxes and physical position along the x-axis show the location of these sequences relative to the Umbonia reference genome. Dashed lines in gold and teal show the expected coverage values for autosomal and X-linked regions, respectively. *Calloconophora* sequences that correspond to the *Umbonia* X chromosome and autosome 9 show reduced coverage consistent with X-linkage.

The presence of an X-autosome fusion in *Calloconophora* would imply a transition from an ancestrally XX/X0 sex chromosome system to a neo-XX/XY system, in which the fused ancestral X + chromosome 9 comprise the neo-X, and the unfused homolog of chromosome 9 forms the neo-Y (Figure 3b). Recombination among sex chromosomes becomes suppressed over time in a stepwise fashion along genomic segments known as evolutionary strata (Charlesworth, et al. 2005; Wright, et al. 2016). Therefore, we examined sex differences in coverage across the neo-X region to ask if strata are present in *Calloconophora* and to gain insight into the relative age of this neo-XX/XY system. This revealed similar levels of sex-differences in coverage among the neo-X and ancestral X (which is entirely hemizygous), suggesting relatively advanced levels of differentiation among neo-X and neo-Y sequences (Figure 3c).

The emergence of a neo-XX/XY system in *Calloconophora* is a marked departure from the long-term X conservation we observe in treehoppers and facilitates an ‘escape’ from the evolutionary trap. Following the autosome-X fusion in *Calloconophora*, the neo-X would have significantly increased its gene content, and a neo-Y would have emerged where there was not one before. The functional impacts of this fusion remain unknown as well as if and how the evolution of these neo-X and neo-Y chromosomes contribute to sex-specific adaptation.

### Testing for X conservation in Auchenorrhyncha

Given that all species in the dataset showed conservation of the ancestral treehopper X chromosome, we compared this ancestral X with the X chromosomes of other members of Auchenorrhyncha (the clade containing treehoppers, leafhoppers, spittlebugs, cicadas, and planthoppers) to investigate sex chromosome evolution across a deeper timescale. We took a similar approach as with the analyses among treehoppers, but this time compared *Umbonia crassicornis* to publicly available chromosome-level assemblies for a leafhopper (Family Cicadellidae; *Homalodisca vitripennis*) and a planthopper (Family Delphacidae; *Nilaparvata lugens*). The divergence time between treehoppers and each group dates to approximately 192 million years ago (mya) and 310 mya, respectively (Johnson, et al. 2018) (Figure 4). In each of the pairwise comparisons, we identified homologous sequences using BLAST (Altschul, et al. 1990) and assessed for concordance in the genomic location of X-linked versus autosomal sequences. In the treehopper to leafhopper comparison, we observed near identical concordance in X-linked versus autosomal regions among the two genomes (Figure 4). When comparing treehoppers with planthoppers, a similar qualitative result was seen, with most treehopper X chromosome sequences also showing X-linkage in the planthopper. Overall, these results indicate conservation of the X chromosome across extremely long time periods spanning the clade Auchenorrhyncha, consistent with recent work describing ancient origins of the insect X chromosome (Chauhan, et al. 2021; Li, et al. 2022; Toups and Vicoso 2023).

**Figure 4.**
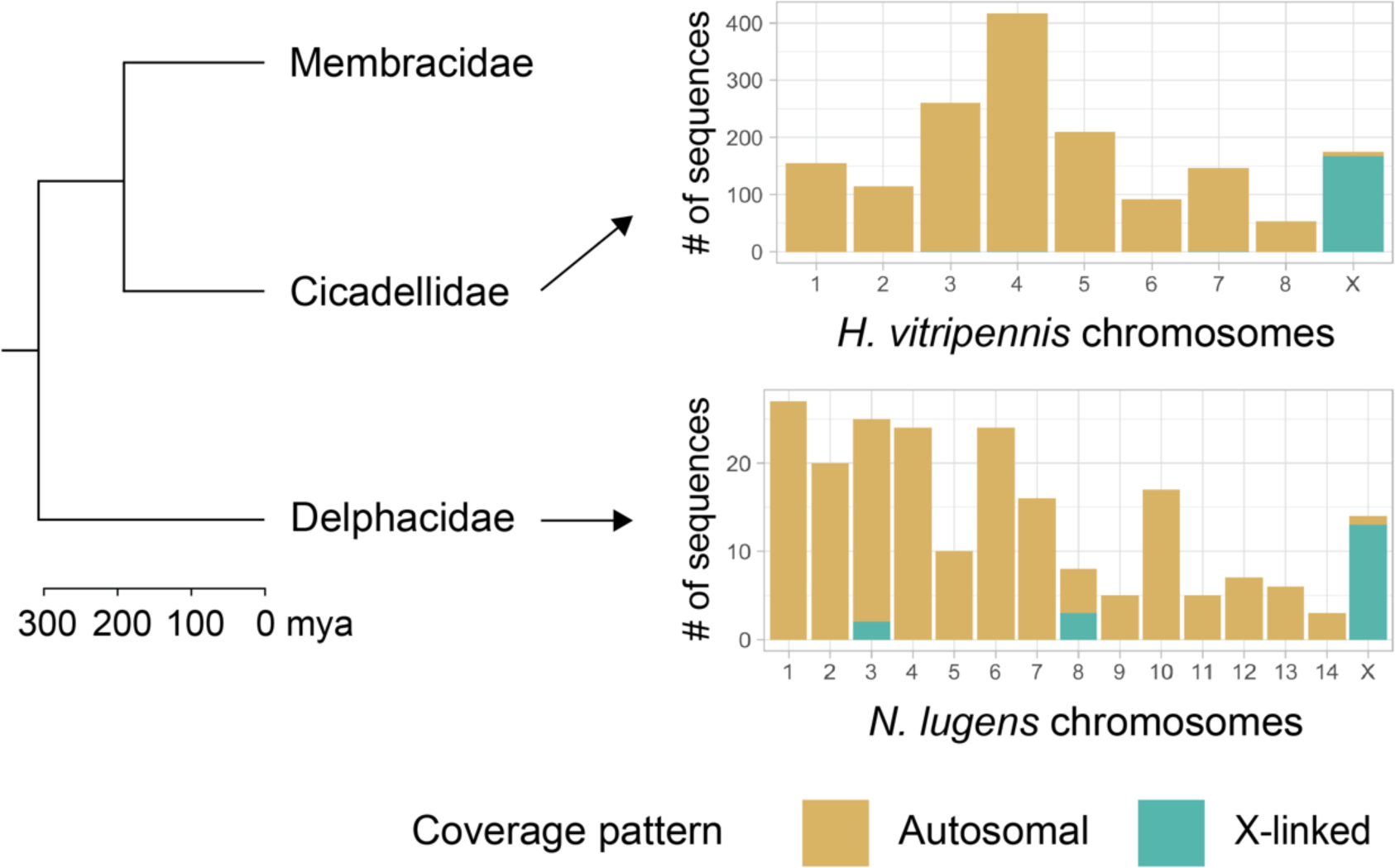
X chromosome identity is conserved within Auchenorrhyncha. Pairwise comparisons between the *Umbonia crassicornis* genome and representative genomes of Cicadellidae (leafhoppers) and Delphacidae (planthoppers). The phylogeny on the left shows the evolutionary distances based on (Johnson, et al. 2018). In the barplots on the right, height of the bar indicates the number of *Umbonia* sequences that correspond to the chromosomes of each species. Color of the bar shows the relative proportion of sequences that are inferred to be autosomal (in gold) versus X-linked (in teal) in *Umbonia*.

Based on published karyotype data, treehoppers exhibit a wide array of chromosomal configurations that suggest frequent genomic rearrangements. The species in our dataset represent a subset of this variation, with chromosome number ranging from 2n=13♂ to 2n=21♂ (Boring 1907; Kornhauser 1914; Halkka 1964; Halkka and Heinonen 1964; Tian and Yuan 1997; Anjos, et al. 2019). The fact that we observed only one species with a chromosomal rearrangement involving a sex chromosome is consistent with recent work in aphids showing conservation of the X between species in spite of extensive autosomal rearrangements (Li, et al. 2020; Mathers, et al. 2021).

Furthermore, it appears that the conservation of the X amidst dynamic autosomal reshuffling extends beyond treehoppers, since we found signatures of shared X identity between treehoppers, leafhoppers, and planthoppers. On the other hand, comparisons within other hemipteran clades like the assassin bugs reveal high levels of synteny across autosomal regions interrupted by recurring X chromosome fission events (Panzera, et al. 1996; Mathers, et al. 2021). Taken together, these results suggest differences among hemipteran clades in their propensity for and/or tolerance of genomic reorganization.

Rearrangements like chromosomal fusions and fissions are largely expected to be deleterious because of their potential to disrupt normal segregation during meiosis (Melters, et al. 2012; Ruckman, et al. 2020). This expectation is based on monocentric organisms (i.e., species with a single, localized centromere per chromosome), in which fusions or fissions can result in chromosome fragments with too many or too few centromeres that fail to segregate properly in meiosis. However, in holocentric species centromeric activity is spread across the chromosome, meaning that fusions and fissions may be better tolerated. Holocentrism has evolved repeatedly in insects (and beyond) and appears to be the ancestral state for treehoppers and the family Hemiptera (Drinnenberg, et al. 2014). The presence of holocentric chromosomes in treehoppers, therefore, may be one of the factors promoting chromosomal evolution in this group, but does not explain why most rearrangements seem to be limited to the autosomes. Further investigation is also needed to understand the apparent differences in rates of chromosomal evolution among holocentric clades.

## Conclusion

We find that X chromosome identity is largely conserved among treehopper species spanning 45 million years. We also find that a chromosomal fusion likely underlies the formation of a neo-XY sex chromosome system. Combining our data with published hemipteran genomes, we observe that the ancestral treehopper X chromosome is homologous to other hemipteran X chromosomes, indicating long-term conservation of the X across more than 300 million years of evolution.

## Materials and Methods

### Sample collection

Adult female and male *Micrutalis calva*, *Thelia bimaculata*, and *Vanduzea arquata* were collected from the wild in New Jersey, USA during July 2019. Adult female and male *Entylia carinata*, *Publilia reticulata*, and *Umbonia crassicornis* were obtained from greenhouse populations housed at Princeton University in September/October 2019.

Adult female and male *Bolbonota melaena*, *Calloconophora caliginosa*, *Cyphonia clavata*, *Cyphonia claviger*, *Membracis foliatafasciata*, *Horiola picta*, and *Neotynelia pubescens* were collected from the campus of UNESP-São Paulo State University, Rio Claro, Brazil between 2020 and 2021.

### Genomic sequencing

For the New Jersey wild-caught samples and Princeton University greenhouse samples, DNA was extracted using a DNeasy Blood and Tissue Kit (Qiagen) following the manufacturer’s protocol. For the samples from Brazil, DNA was extracted using a Wizard Genomic DNA purification kit (Promega, WI, USA). All libraries were prepared and sequenced at the Center for Genomic Research at the University of Liverpool using standard protocols. DNA was sequenced on the Illumina NovaSeq 6000 S1 and S4, resulting in on average 259 million 150bp paired-end reads per individual.

### Umbonia crassicornis genome assembly

#### Genome assembly

The genome of *Umbonia crassicornis* was assembled using a single lab-reared adult female (collected December 2016) that originated from a population in Ft. Lauderdale, FL. Linked-read libraries were constructed using the 10x Genomics platform and sequenced on the Illumina HiSeq X platform, generating 150 base paired-end reads from approximately half a lane. An additional lab-reared female from the same host plant was collected for Hi-C library preparation to aid in scaffolding the genome assembly.

#### Laboratory methods

High molecular weight (HMW) DNA was extracted using Qiagen Genomic Tip kits (Qiagen, USA, Catalog #10223) with slight modifications to the manufacturer’s protocol. Specimens were gently homogenized with a pestle over dry ice, followed by the addition of 350 ul of buffer ATL and 4ul of RNAse A. After incubation at 37C for 30 minutes, 50ul of proteinase K and 1 mL of G2 buffer were added, and the samples were incubated overnight at 50 C. The standard genomic tip protocol was then followed, with centrifugations performed at 12,000g for 30 and 15 minutes. DNA was eluted in 50ul of TE buffer.

#### HiC sequencing

To scaffold the 10X genome drafts, *in situ* Hi-C libraries were prepared following a previously described method (Jones, et al. 2023). Tissue from a single individual was crosslinked, and nuclei were lysed while maintaining their integrity. DNA was restricted, and overhangs were filled in with a biotinylated base. Free ends were then ligated together *in situ*, followed by reversal of crosslinks and shearing of DNA to 300-500bp fragments. Biotinylated ligation junctions were isolated using streptavidin beads, and the recovered material was used for Illumina library construction. This involved end-repair using T4 DNA polymerase, Klenow polymerase, and T4 polynucleotide kinase, followed by A-tailing with Klenow fragment (3’ to 5’ exo minus) and dATP. Illumina adapters with a single ‘T’ base overhang were ligated to the DNA fragments. The ligated DNA was PCR amplified for 8-12 cycles using Illumina primers, and library fragments of 400-600 bp were purified using SPRI beads. The purified DNA was captured on an Illumina flow cell for cluster generation and sequenced following the manufacturer’s protocols.

### Identification and comparison of X-linked sequences

For *Umbonia crassicornis*, paired-end reads from two males and two females were mapped to the reference genome using BWA (Li and Durbin 2009) version 0.7.17 with default settings. Uniquely mapped reads were extracted using the grep command “XT:A:U”. SOAPcov v2.7.9 (https://github.com/aquaskyline/SOAPcoverage) was then used to calculate coverage depth for each scaffold, and in 50kb windows across the genome. The log ratio of male to female coverage was calculated for each scaffold, and for each window, using a custom script.

For each of the remaining species, female paired-end reads were used for *de novo* genome assembly using SOAPdenovo2 (Luo, et al. 2012) r242. All reads were used during the contig and scaffold assembly steps and option -F was used during scaffolding. We used GapCloser version 1.12 (https://anaconda.org/bioconda/soapdenovo2-gapcloser) to close gaps generated in the scaffolding step. The optimum kmer value for each assembly was determined using kmergenie (Chikhi and Medvedev 2014) version 1.7051. Once each species’ genome had been assembled, we mapped the male and female reads of each species to its genome assembly using BWA (Li and Durbin 2009) version 0.7.17 with default settings. We then used SOAPcov v2.7.9 (https://github.com/aquaskyline/SOAPcoverage) to calculate the coverage depth for each scaffold and used our custom script to obtain the log ratio of male to female coverage. Scaffolds with log_2_(M:F) coverage less than the median coverage – 0.5 were designated as X-linked. Scaffolds with log_2_(M:F) coverage greater than or equal to the median coverage – 0.5 were designated as autosomal.

To compare X chromosome identity among taxa, we used BLAST to assign inferred coding sequences from each taxon to a chromosomal location in the *Umbonia* genome. We first downloaded a publicly available transcriptome assembly for *Entylia carinata* (Fisher, et al. 2020). These sequences were BLASTed against the *Umbonia crassicornis* genome assembly using blastn (Altschul, et al. 1990) version 2.11.0+ with parameters -perc_identity 30 -evalue 10e-10. In cases where there was more than one hit, the match with the higher bitscore and percent identity was chosen. We then BLASTed the *E. carinata* transcriptome sequences to each of our *de novo* assemblies, again using parameters -perc_identity 30 -evalue 10e-10 and choosing the match with the higher bitscore and percent identity in cases of multiple hits. Next, for each taxon we intersected these BLAST results and the log_2_(M:F) coverage values using a custom script to assign putative autosomal and X-linked sequences to their corresponding location in the *U. crassicornis* genome.

### Comparing X-linked sequences among Auchenorrhynchan families

We downloaded the coding sequences and gtf/gff annotation files for two previously published chromosome-level genome assemblies of representative Auchenorrhynchan species: *Homalodisca vitripennis* (Family Cicadellidae), and *Nilaparvata lugens* (Family Delphacidae).

The *H. vitripennis* data (UT_GWSS_2.1) were downloaded from NCBI (https://www.ncbi.nlm.nih.gov/datasets/genome/GCF_021130785.1/), and the *N. lugens* data were downloaded from InsectBase (http://v2.insect-genome.com/Organism/572). For each species, coding sequences were BLASTed against the *E. carinata* transcriptome (Fisher, et al. 2020) using parameters -perc_identity 30 -evalue 10e-10 and choosing the match with the higher bitscore and percent identity in cases of multiple hits. We then used each genome’s annotation file to locate X-linked and autosomal sequences and compared these to our previous BLAST results between *E. carinata* and *U. crassicornis* to determine whether they correspond to X-linked or autosomal treehopper sequences.

### Cytology

Chromosomal analyses were performed by inspection of meiotic and mitotic cells obtained from male testis stained with DAPI (4′,6-diamidino-2-phenylindole). The diploid number and sex chromosomes were determined here for *C. claviger*, *C. clavata*, and *C. caliginosa*, while for the other species the data are published (Anjos, et al. 2019), except for *P. reticulata* in which we did not have proper material for chromosomal analysis (Supplementary Table 1). For the fluorescent *in situ* (FISH) mapping of telomeric probe in *C. caliginosa* we followed a published protocol (Cabral-de-Mello and Marec 2021). The insect telomeric probe was synthesized by non-template PCR according to (Ijdo, et al. 1991) using the self-complementary primers (TTAGG)_5_ and (CCTAA)_5_ and labeled with digoxigenin-11-dUTP by nick translation.

## Acknowledgements

This work was funded by a NERC Independent Research Fellowship to A.E.W. (NE/N013948/1), and an NSF PRFB (DBI-1812164) and Michigan State University Presidential Postdoctoral Fellowship to D.H.P.D. S.D.K. is an HHMI Freeman Hrabowski Scholar. M.P.F. was supported by an NIH training grant through the Lewis-Sigler Institute for Integrative Genomics. We thank the Slate lab and Wright lab for helpful comments on the analyses and manuscript and Chung-Ping Lin for help with determining the species identification for the *Calloconophora* samples. The laboratory work was supported by the UK Natural Environment Research Council (NERC) Environmental Omics Facility. Data generation was carried out by the Centre for Genomic Research, which is based at the University of Liverpool. We acknowledge IT Services at The University of Sheffield and the Texas Advanced Computing Center (TACC) at The University of Texas at Austin for providing HPC resources that have contributed to the research results reported within this paper.

## Data Accessibility

All raw sequence data are deposited under BioProject ID PRJNA1106845 and the *Umbonia crassicornis* genome assembly is deposited under BioProject ID PRJNA1122077 in the NCBI Sequence database.

## Notes

### Competing Interest Statement

The authors have declared no competing interest.

