## Supplementary Tables for "Chromosomal fusion drives sex chromosome evolution in treehoppers despite long-term X chromosome conservation"

**Supplementary Table 1. Sample, karyotype, and sequencing information for taxa included in this study.**

| Sample | Species | Sex | Sex chromosome system | Chromosome number (female/male) | Total # of Reads | Total # bases | Approx coverage based on 1.2Gb genome |
|---|---|---|---|---|---|---|---|
| DP1 | *Entylia carinata* | F | X0 | 22/21 | 236013750 | 17701031250 | 14.75 |
| DP2 | *Entylia carinata* | M | X0 | 22/21 | 240378162 | 18028362150 | 15.02 |
| DP3 | *Entylia carinata* | F | X0 | 22/21 | 229378952 | 17203421400 | 14.34 |
| DP4 | *Entylia carinata* | M | X0 | 22/21 | 211402100 | 15855157500 | 13.21 |
| DP5 | *Micrutalis calva* | F | X0 | 22/21 | 259345838 | 19450937850 | 16.21 |
| DP6 | *Micrutalis calva* | M | X0 | 22/21 | 232935414 | 17470156050 | 14.56 |
| DP7 | *Micrutalis calva* | F | X0 | 22/21 | 211697338 | 15877300350 | 13.23 |
| DP8 | *Micrutalis calva* | M | X0 | 22/21 | 286085784 | 21456433800 | 17.88 |
| DP9 | *Publilia reticulata* | F | X0 | ? | 230554556 | 17291591700 | 14.41 |
| DP10 | *Publilia reticulata* | M | X0 | ? | 190511034 | 14288327550 | 11.91 |
| DP11 | *Publilia reticulata* | F | X0 | ? | 235410126 | 17655759450 | 14.71 |
| DP12 | *Publilia reticulata* | M | X0 | ? | 229894648 | 17242098600 | 14.37 |
| DP13 | *Thelia bimaculata* | F | X0 | 22/21 | 262178548 | 19663391100 | 16.39 |
| DP14 | *Thelia bimaculata* | M | X0 | 22/21 | 247517894 | 18563842050 | 15.47 |
| DP15 | *Thelia bimaculata* | F | X0 | 22/21 | 287138902 | 21535417650 | 17.95 |
| DP16 | *Thelia bimaculata* | M | X0 | 22/21 | 239004780 | 17925358500 | 14.94 |
| DP17 | *Umbonia crassicornis* | F | X0 | 20/19 | 217438244 | 16307868300 | 13.59 |
| DP18 | *Umbonia crassicornis* | M | X0 | 20/19 | 211628288 | 15872121600 | 13.23 |
| DP19 | *Umbonia crassicornis* | F | X0 | 20/19 | 217901640 | 16342623000 | 13.62 |
| DP20 | *Umbonia crassicornis* | M | X0 | 20/19 | 121994174 | 9149563050 | 7.62 |
| DP21 | *Vanduzea arquata* | F | X0 | 18/17 | 363207514 | 27240563550 | 22.70 |
| DP22 | *Vanduzea arquata* | M | X0 | 18/17 | 331431040 | 24857328000 | 20.71 |
| DP23 | *Vanduzea arquata* | F | X0 | 18/17 | 233798694 | 17534902050 | 14.61 |
| DP24 | *Vanduzea arquata* | M | X0 | 18/17 | 269454890 | 20209116750 | 16.84 |
| 1C | *Horiola picta* | F | X0 | 22/21 | 214832248 | 16112418600 | 13.43 |
| 1D | *Horiola picta* | F | X0 | 22/21 | 214542278 | 16090670850 | 13.41 |
| 2B | *Horiola picta* | M | X0 | 22/21 | 164428916 | 12332168700 | 10.28 |
| 2D | *Horiola picta* | M | X0 | 22/21 | 235295432 | 17647157400 | 14.71 |

| | | | | | | | |
|---|---|---|---|---|---|---|---|
| 3B | *Neotynelia pubescens* | F | X0 | 22/21 | 206263678 | 15469775850 | 12.89 |
| 3D | *Neotynelia pubescens* | F | X0 | 22/21 | 219214080 | 16441056000 | 13.70 |
| 4B | *Neotynelia pubescens* | M | X0 | 22/21 | 197045214 | 14778391050 | 12.32 |
| 4D | *Neotynelia pubescens* | M | X0 | 22/21 | 296574192 | 22243064400 | 18.54 |
| 5B | *Membracis foliatafasciata* | F | X0 | 14/13 | 150032424 | 11252431800 | 9.38 |
| 5D | *Membracis foliatafasciata* | F | X0 | 14/13 | 256359460 | 19226959500 | 16.02 |
| 6B | *Membracis foliatafasciata* | M | X0 | 14/13 | 180297230 | 13522292250 | 11.27 |
| 6D | *Membracis foliatafasciata* | M | X0 | 14/13 | 278931886 | 20919891450 | 17.43 |
| 9B | *Cyphonia claviger* | F | X0 | 20/19 | 222563856 | 16692289200 | 13.91 |
| 10B | *Cyphonia claviger* | F | X0 | 20/19 | 436910698 | 32768302350 | 27.31 |
| 12B | *Cyphonia claviger* | M | X0 | 20/19 | 149390512 | 11204288400 | 9.34 |
| 14B | *Cyphonia clavata* | F | X0 | 20/19 | 188388296 | 14129122200 | 11.77 |
| 7C | *Cyphonia clavata* | M | X0 | 20/19 | 248655672 | 18649175400 | 15.54 |
| R10 | *Cyphonia clavata* | M | X0 | 20/19 | 1121509792 | 84113234400 | 70.09 |
| R6 | *Cyphonia clavata* | M | X0 | 20/19 | 1251273410 | 93845505750 | 78.20 |
| 17B | *Bolbonota melaena* | F | X0 | 22/21 | 204807698 | 15360577350 | 12.80 |
| 10C | *Bolbonota melaena* | F | X0 | 22/21 | 248310210 | 18623265750 | 15.52 |
| 11C | *Bolbonota melaena* | F | X0 | 22/21 | 211829260 | 15887194500 | 13.24 |
| 20B | *Bolbonota melaena* | M | X0 | 22/21 | 151330058 | 11349754350 | 9.46 |
| 7D | *Calloconophora caliginosa* | F | XY | 18/18 | 280988320 | 21074124000 | 17.56 |
| 8D | *Calloconophora caliginosa* | M | XY | 18/18 | 250061914 | 18754643550 | 15.63 |

**Supplementary Table 2. Assembly statistics for *de novo* genome assemblies from Illumina short reads**

| | Number of sequences | Average length | Maximum length | Total length | N50 |
|---|---|---|---|---|---|
| *Bolbonota melaena* | 7,891,336 | 202 | 11612 | 1,596,938,007 | 226 |
| *Calloconophora caliginosa* | 5,881,810 | 312 | 49683 | 1,833,318,984 | 492 |
| *Cyphonia clavata* | 4,755,768 | 223 | 3872 | 1,059,558,444 | 268 |
| *Cyphonia claviger* | 9,895,077 | 194 | 5443 | 1,923,787,491 | 210 |
| *Entylia carinata* | 1,736,639 | 359 | 11903 | 624,140,518 | 604 |
| *Horiola picta* | 6,276,945 | 287 | 20878 | 1,799,441,780 | 592 |
| *Membracis foliatafasciata* | 3,839,033 | 238 | 7888 | 913,223,601 | 333 |
| *Micrutalis calva* | 9,642,625 | 283 | 140306 | 2,730,412,327 | 426 |
| *Neotynelia pubescens* | 3,352,327 | 192 | 3652 | 643,739,419 | 204 |
| *Publilia reticulata* | 4,050,913 | 431 | 41981 | 1,745,737,668 | 1290 |
| *Thelia bimaculata* | 5,944,557 | 326 | 44828 | 1,935,543,686 | 628 |
| *Vanduzea arquata* | 4,821,435 | 325 | 39852 | 1,567,769,002 | 720 |
