## Supplementary Figure 1 for "Chromosomal fusion drives sex chromosome evolution in treehoppers despite long-term X chromosome conservation"

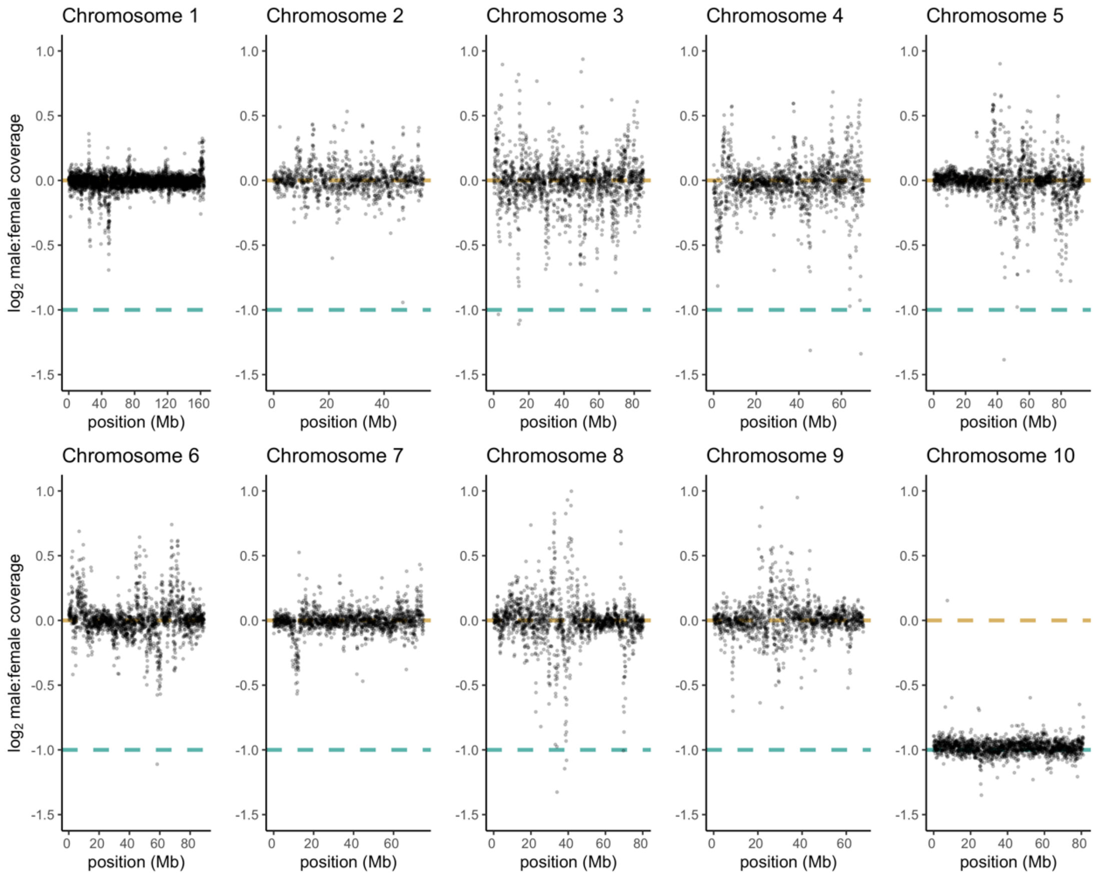


**Supplementary Figure 1. Sex differences in coverage across *Umbonia crassicornis* chromosomes show a consistent reduction of male coverage across the length of chromosome 10.** Dots show male/female coverage ratio in 50kb windows. Dashed lines in gold and teal show the expected coverage values for autosomal and X-linked regions, respectively. Chromosome 10 has reduced male coverage across its entire length, consistent with an XX/X0 sex chromosome system.
